# Microbiome depletion and recovery in the sea anemone, *Exaiptasia diaphana*, following antibiotic exposure

**DOI:** 10.1101/2023.12.13.571442

**Authors:** Sophie MacVittie, Saam Doroodian, Aaron Alberto, Maggie Sogin

## Abstract

Microbial species that comprise host-associated microbiomes play an essential role in maintaining and mediating the health of plants and animals. While defining the role of individual or even complex communities is important towards quantifying the effect of the microbiome on host health, it is often challenging to develop causal studies that link microbial populations to changes in host fitness. Here, we investigated the impacts of reduced microbial load following antibiotic exposure on the fitness of the anemone, *Exaiptasia diaphana* and subsequent recovery of the host’s microbiome. Anemones were exposed to two different types of antibiotic solutions for three weeks and subsequently held in sterilized seawater for a subsequent three-week recovery period. Our results revealed that both antibiotic treatments reduced the overall microbial load during and up to one week post treatment. The observed reduction in microbial load was coupled to reduced anemone biomass, halted asexual reproduction rates, and for one of the antibiotic treatments, the partial removal of the anemone’s algal symbiont. Finally, our amplicon sequencing results of the 16S rRNA gene revealed that anemone bacterial composition only shifted in treated individuals during the recovery phase of the experiment, where we also observed a significant reduction in the overall diversity of the microbial community. Our work implies that the *E. diaphana’s* microbiome contributes to host fitness and that the recovery of the of the host’s microbiome following disturbance with antibiotics leads to a reduced, but stable microbial state.

**Importance:** *Exaiptasia diaphana* is an emerging model used to define the cellular and molecular mechanisms of coral-algal symbioses. *E. diaphana* also houses a diverse microbiome, consisting of hundreds of microbial partners with undefined function. Here, we applied antibiotics to quantify the impact of microbiome removal on host fitness as well as define trajectories in microbiome recovery following disturbance. We showed that reduction of the microbiome leads to negative impacts on host fitness, and that the microbiome does not recover to its original composition while held under aseptic conditions. Rather the microbiome becomes less diverse, but more consistent across individuals. Our work is important because it suggests that anemone microbiomes play a role in maintaining host fitness, that they are susceptible to disturbance events, and it is possible to generate gnotobiotic individuals that can be leveraged in microbiome manipulation studies to investigate the role of individual species on host health.

## Introduction

Most animals rely on a complex microbiome to support individual fitness, heath, and metabolism (1, 2). For example, microbiomes provide animals with essential nutrients, support reproductive pathways and protect hosts from disease causing pathogens and toxic compounds (as reviewed in (3)). An emerging hypothesis in microbiome research is that the unit of selection is indeed the “metaorganism”, which is defined as the animal host together with its archaeal, bacterial, fungal, viral and microeukaryote associates (4). One challenge in understanding the role of individual microbial partners within the metaorganism is that for many systems we lack causal studies that support the intrinsic role of the microbiome in determining the phenotype of the host (but see (5–7)).

Marine cnidarians, such as jellyfish, sea anemones, and reef-building corals, host a broad diversity of microbial species within their tissue layers (8–11). Some of these microbial taxa are thought to play a fundamental role in supporting host health and metabolism. The most famous of which is the relationship between reef-building corals and their dinoflagellate, microalgal symbionts in the family *Symbiodiniaceae*. The microalgal partners support coral growth and survival through the transfer of sugars to the animal host. In addition to *Symbiodiniaceae*, many cnidarians also require other microbial species (e.g., bacteria and fungi), to support complex metabolic pathways within the metaorganism (12). For example, cnidarian associated bacteria are likely involved in nitrogen and sulfur metabolism, and nutrient cycling (13–18), defense against host pathogens (5, 19–21), and, therefore, intrinsically supports coral health (e.g., 19, 22, 23). Understanding the individual and collective role of these microbes in supporting cnidarians is critical as researchers and conservationists alike aim to develop new approaches (e.g., beneficial microorganisms for corals) for mitigating the impacts of environmental stress on coral reefs (3, 24, 25). Yet, much of the current work in defining microbial function of cnidarians remains correlative as there is a need to develop reductionist approaches that allow for the quantification of the impacts of individual microbial species or consortia on host health, metabolism, and fitness (26).

One promising approach in defining the role of individual microbes within metaorganisms is to rear hosts in the absence of their microbiome or with a reduced microbial load (e.g., 5, 27). By generating these so-called gnotobiotic (known suite of microbial partners), or axenic (germ-free) systems, we can begin to dissect the individual roles of microbial species within the metaorganism and their impacts on cnidarian health. For example, the depletion of *Hydra’s* microbiome revealed that microbial taxa were (i) critical in defending the anemone against fungal pathogens (5), (ii) involved in cell signaling pathways that controlled host development (28), and (iii) regulated host physiology and phenotype (29, 30). In marine cnidarians, most of the microbiome depletion work is focused on quantifying the impacts of *Symbiodiniaceae* after removal from host tissues, however other studies are beginning to quantify the reduction of other microbial species within the metaorganism (7, 27, 31–33). Here, we aimed to expand on these initial studies by (1) quantifying the impact of reducing the microbiome on cnidarian physiology and (2) documenting patterns in microbiome recovery following disturbance using a longitudinal sampling approach. By quantifying shifts in cnidarian microbiomes through longitudinal sampling, we aim to describe community dynamics related to the recovery of cnidarian microbiomes. Defining how the microbiome recovers following disturbance is critical when assessing the efficacy and practicality of probiotic approaches in marine habitats. To meet these aims, we conducted a microbiome manipulation experiment using the sea anemone, *Exaiptasia diaphana*, hereon referred to as Aiptasia.

Aiptasia is an emerging model system for exploring the role of microbial partners within marine cnidarians (34). Aiptasia houses the same type of *Symbiodiniaceae* partners within its gastrodermal tissue layer as reef building corals. Consequently, much of the current work in Aiptasia is focused on studying shifts in the cell and molecular machinery following the removal of the algal symbiont (i.e., bleaching)(35–37). However, like corals, Aiptasia also hosts a complex microbiome consisting of approximately 100 bacterial species that live in the anemone’s tissue and mucus layers (10, 32, 38, 39). It is possible to reduce the microbial load of the Aiptasia microbiome using antibiotics (31), however microbial depletion is unstable in the presence of biofilms (32), and thus it is difficult to maintain in anemone cultures. Importantly, it is not clear how the depletion of the microbiome directly impacts overall Aiptasia physiology and health. To begin to disentangle the roles of individual microbes within Aiptasia, it is essential to quantify how the microbial community shifts during depletion and what the overall impact is on the metaorganism.

In this study, we set out to quantify the effects of antibiotic treatment on the bacterial load and composition of the Aiptasia microbiome while testing to determine if our experimental treatments impacted metaorganism fitness. Leveraging lessons learned from past knockdown studies (31, 32), we exposed individual sea anemones to two different antibiotic solutions for three weeks and monitored the impact on the microbiome (i.e., both alpha and beta diversity) and physiology (i.e., total biomass, algal cell density, and asexual reproduction rate) during recovery. Overall, our results revealed that exposure to antibiotics reduced the abundance of the Aiptasia bacterial community, shifted the composition of community members, which resulted in fitness declines across measured metrics.

## Results

### Experimental Overview

We set out to quantify the impacts of antibiotic exposure on the disturbance and recovery of the Aiptasia (host strain H2 with *Breviolum minutum*) microbiome and associated host fitness. The 76-day long experiment was divided into three phases: priming, treatment, and recovery (**Fig. 1A**). During the priming, anemones that were treated with antibiotic solutions during the treatment phase were first held in filtered artificial seawater (FASW) for 33 days to reduce overall microbial load. After which, these anemones were treated with either antibiotic solution 1 (ABS1; 50µg/ml of carbenicillin, chloramphenicol, nalidixic acid and rifampicin) or antibiotic solution 2 (ABS2; 50µg/ml of neomycin, penicillin, rifampicin, streptomycin) for a total of 22 days. During recovery, treated anemones were held in FASW for 21 days. Throughout the experiment, we sampled individual anemones from the treatment and control groups (i.e., individuals held in artificial seawater (ASW)) on days 0 (baseline), 33 (priming), 55 (treatment), 61 (recovery 1) and 76 (recovery 2) for amplicon sequencing using the V4 region of the 16S SSU of the rRNA gene, bacterial load, biomass, *Symbiodiniaceae* density and asexual reproduction rate (**Fig. 1A**).

**Figure 1.**
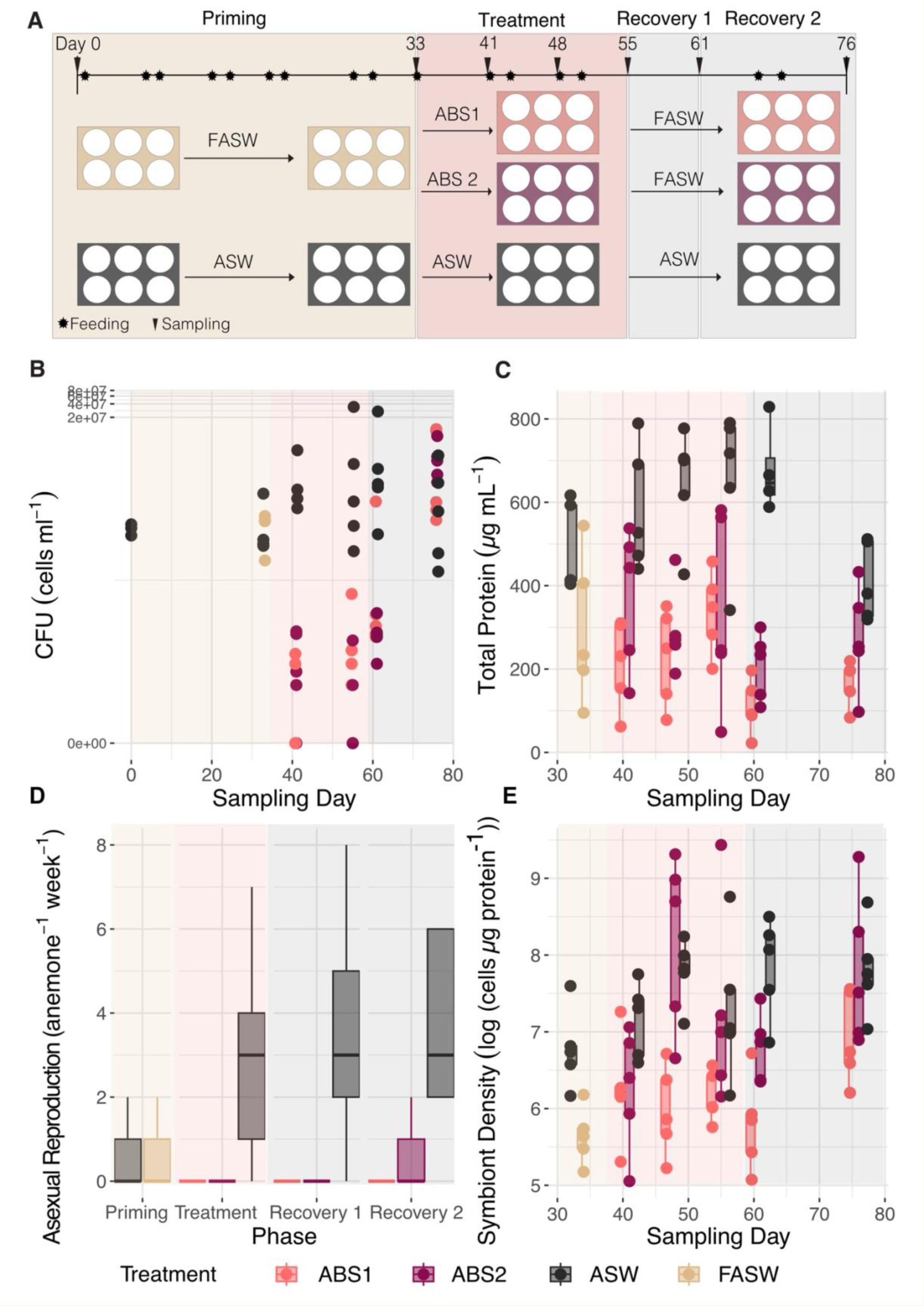
Antibiotic exposure reduced bacterial load and host fitness. **A)** Experimental overview outlining the steps in the 76-day long experiment. First, individual Aiptasia polyps were held in either artificial seawater (ASW; control) or filtered artificial seawater (FASW) for 33 days. Anemones in FASW were then treated with either ABS1 or ABS2 solutions for 22 days and subsequently allowed to in FASW during for 21 days. Anemones were sampled (black triangle) at least 4 days post feeding to avoid sampling shifts in microbiomes related to heterotrophic feeding (black star). **B)** Colony forming unit counts (CFUs, cells mL^-1^) significantly differed across treatments and sampled time points (2-way ANOVA *p-value* < 0.001). Aiptasia exposed to antibiotics experienced a significant (Tukey HSD adjusted *p-value* < 0.001) reduction in bacterial load during treatment on day 41 and 55, and during the first phase of recovery on day 61. CFU counts returned to control levels on day 76(day 0 n=3, otherwise n=5; day 48 CFU counts were excluded due to plate contamination). **C)** Total protein concentrations differed significantly across time (2-way ANOVA *p-value* = 0.01) and experimental treatments (2-way ANOVA *p-value* < 0.001) between treated anemones and the control group. Boxplots of total protein (µg mL^-1^) measured from each polyp (n=5), revealed that anemone biomass significantly declined (Tukey HSD *adjusted p-value* < 0.001) initially during the priming phase when individuals were held in FASW in comparison to the control and remained significantly (Tukey HSD *adjusted p-value* p < 0.001) reduced in concentration in comparison to the control throughout the remainder of the experiment. **D)** Exposure to antibiotics significantly reduced (Kruskal-Wallis p-value < 0.001) the asexual reproduction rate of individual anemones (n=6) during treatment and recovery. Boxplots of asexual reproduction rate as measured by number of observed pedal lacerates per individual per week showed that both antibiotics halted pedal laceration rate until the final recovery time point, when anemones held in ABS2 started to recovery their capacity to undergo asexual reproduction. **E)** Algal cell densities (cell density µg protein^-1^) differed significantly across time (2-way ANOVA *p-value* < 0.001) and experimental treatments (2-way ANOVA *p-value* < 0.001) between treated anemones and the control group(n=5). Boxplots of normalized symbiont density show that FASW reduced algal cell density during the priming phase. While anemones in ABS2 recovered their algal populations during treatment, anemones held in ABS1 had reduced algal population until the final recovery period. See **Table S1** for all model statistics and **Table S2** for means and standard errors.

### Exposure to antibiotics depletes Aipstasia’s bacterial load during treatment

Anemones treated with both ABS1 and ABS2 solutions had a significantly reduced bacterial load, as measured by colony forming unit counts (CFU), during treatment and up to one-week of recovery in FASW (Kruskal-Wallis p-value < 0.001; **Fig. 1B; Table S1, S2**). Unlike previous studies (32, 40), there were no differences (p > 0.05) in CFU counts between anemones held in FASW (average CFU cells / mL = 8.1×10^4^ ± 2.3×10^4^) in comparison to the controls (average CFU cells / mL= 1.2×10^5^ ± 8.9×10^4^) during priming. Rather, CFU counts only significantly decreased (p < 0.05) in anemones exposed to the ABS solutions during treatment (day 41 and 55; average CFU cells / mL = 32 ± 2.5 - 504 ± 4.2) in comparison to the controls (average CFU cells / mL = 1.1×10^6^ ± 8.3×10^5^ - 7.2×10^6^ ± 6.7×10^6^). During recovery in FASW, CFU counts remained significantly lower (p < 0.05) in treated anemones for up to one week post treatment (day 61; average CFU cells / mL = 336 ± 12 - 5.2×10^4^ ± 5.1×10^4^) in comparison to the control (average CFU cells / mL = 5.9×10^6^ ± 5.2×10^5^). However, by three weeks post treatment (day 76) CFU counts returned to similar levels as the control group (average CFU cells / mL = 7.3×10^5^ ± 5.3×10^5^ - 4.2×10^6^ ± 2.5×10^6^). Throughout the experiment, there were no significant differences (p > 0.05) in CFU counts between anemones held in ABS1 versus ABS2 solutions (**Table S1**).

### Exposure to antibiotics reduced anemone fitness

Anemones treated with ABS1 and ABS2 had reduced total protein content in comparison to the controls during treatment and recovery, suggesting our treatment regimes led to significant decreases in overall biomass (**Fig. 1C; Tables S1, S2**). While total protein content in anemones held in FASW (average µg/ml of protein = 295.13 ± 79.90) was decreased in comparison to the control (average µg/ml of protein = 487.95±47.94), differences between treatments was not significant. Total protein content was significantly reduced (*p-value* < 0.01) after only seven days of exposure in ABS1 (average µg/ml of protein = 212.60 ± 47.25) compared to the control (average µg/ml of protein = 583.98 ± 67.06), whereas ABS2 was slightly reduced but not significantly different from the control (average µg/ml of protein = 372.01 ± 76.09). Total protein content was significantly (*p-value* <0.05) reduced in ABS1 and ABS2 throughout the remainder of treatment (**Tables S1, S2**). During recovery at day 61, anemones continued to diminish in size in both ABS1 (average µg/ml of protein = 109.46±29.53) and ABS2 (average µg/ml of protein = 206.86 ± 36.02) while the control maintained their size (average µg/ml of protein = 677.62 ± 52.78). At the final recovery time point (76 d), treated anemones began to increase in biomass (average µg/ml of protein = 168.36 ± 24.31 - 274.79 ± 56.18), which was no longer significantly different from the control (average µg/ml of protein = 409.12 ± 42.06).

Anemones treated with antibiotics had reduced asexual reproduction rate, as measured by pedal laceration, during treatment (Kruskal-Wallis test, p < 0.0001; **Fig. 1D, Table S1, S2**). FASW treatment during priming did not impact on the rate of pedal laceration (*p-value* > 0.05; mean pedal laceration rate = 0.26 ± 0.11 individuals week^-1^). During recovery, anemones held in ABS1 did not pedal lacerate even during the final sampling time point, while anemones held in ABS2 began to pedal lacerate at 71 d (mean pedal laceration rate = 0.60 ± 0.40 individuals week^-1^). Both rates were significantly lower than the control (mean pedal laceration rate = 3.5 ± 0.92 individuals week^-1^).

Symbiont density did not vary significantly in individuals treated with ABS2 or FASW (2- way ANOVA, *p-value* > 0.1, **Fig. 1E, Table S1, S2**). In contrast, symbiont density decreased in anemones treated with ABS1 during treatment and in the first week of recovery. Two weeks into treatment (day 48), symbiont density in ABS1 (average cell/µg protein = 447.10 ± 114.63) was significantly lower (*p-value* < 0.05) compared to the control (average cell/µg protein = 2570.09 ± 419.78). After one-week of recovery, anemones in ABS1 were visually lighter and had a significantly lower (p < 0.01) symbiont density (average cell/µg protein = 388.21 ± 117.10) when compared to the control (average cell/µg protein = 2963.21 ± 700.96). By the end of the recovery period, symbiont density in ABS1 (average cell/µg protein = 1167.09 ± 297.94) began to return to control levels (average cell/µg protein = 2851.40 ± 814.97). While there were no discernible patterns in ABS2 samples, Aiptasia exposed to ABS2 exposed had much higher variability in protein content than all other treatments throughout the experiment.

### Exposure to antibiotics shifts the composition of the core Aiptasia microbiome

To quantify the impacts of antibiotic exposure on the Aiptasia microbiome, we sequenced the V4 region of the 16S rRNA gene from individual anemones. In total, we recovered between 83 and 890,282 reads from each sample. Samples and extraction blanks with fewer than 20,000 or greater than 250,000 reads were removed from the analysis. Removed samples included four individuals treated with ABS solutions, one control sample, two filtered seawater samples and one extraction blank. After quality filtering the read set, determining amplicon sequence variants (ASVs), and removal of contaminating, ribosomal, and chromosomal ASVs, the resulting dataset contained 412 individual taxa across 82 samples. To determine how antibiotics impacted the Aiptasia microbiome, we first compared alpha diversity across treatments and sampling time points (day 0, 33, 55,61 and 76; **Fig. 2A**). We then calculated the core (**Fig. 2D**) membership based on prevalence and abundance of ASVs from the control samples. Finally, we assessed shifts in beta-diversity of the core (**Figs. 2B, C**) across treatments and sampling time points (day 0, 33, 55,61 and 76).

**Figure 2.**
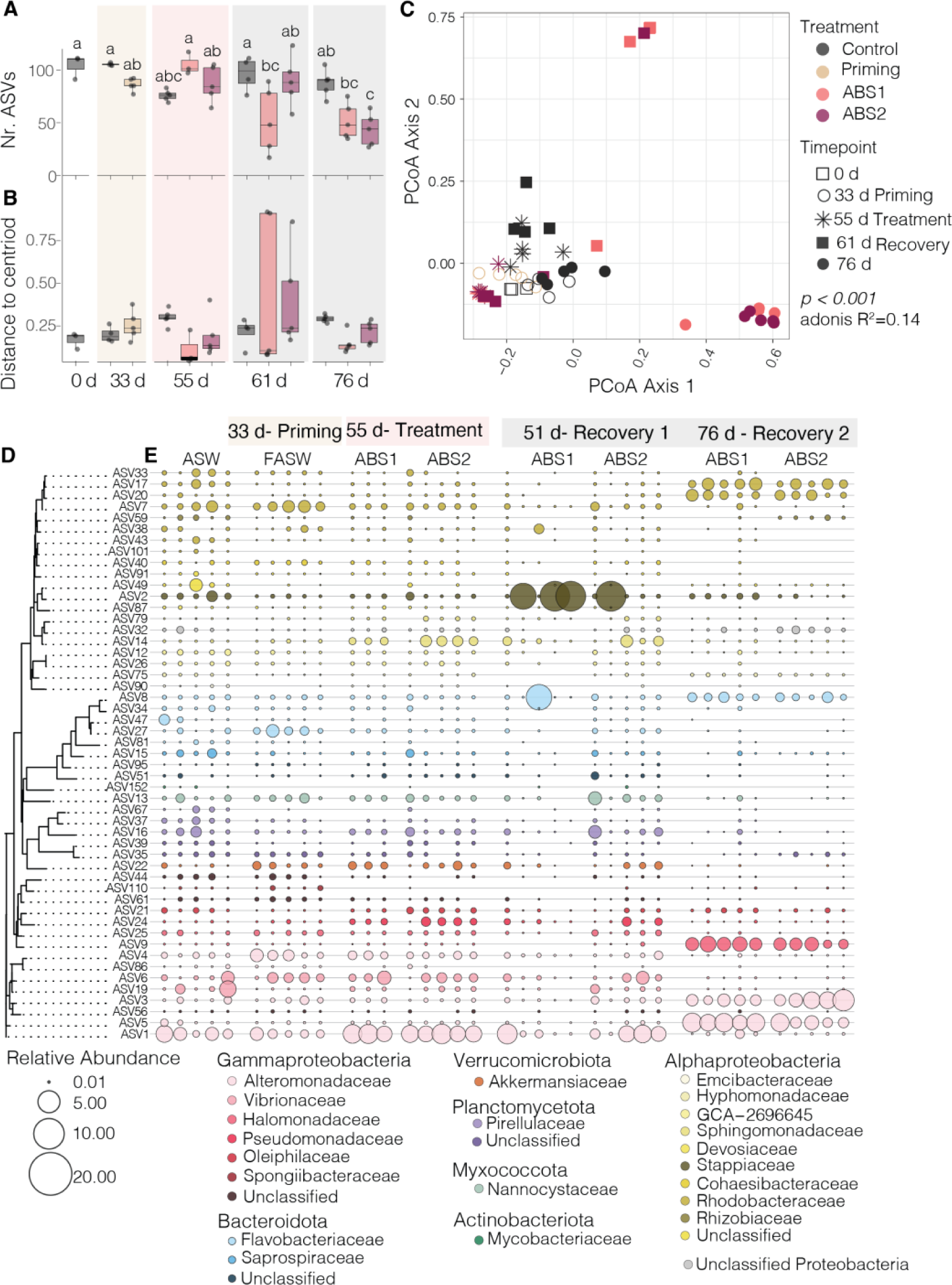
Antibiotic exposure shifts the core microbiome of Aiptasia. **A)** Boxplots showing alpha diversity, as reported as the number of observed ASVs, revealed that anemones exposed to antibiotics had a significantly (two-way ANOVA *p-value* < 0.001) less diverse community in comparison to the controls (n=4-5 per treatment per time point). **B)** Boxplots of beta-dispersion values and **C)** a PCoA analysis calculated from a Bray-Curtis distance matrix of the core (**D**) microbiome revealed that beta-diversity significantly differs (ANOSIMS R^2^=0.14, *p-value* < 0.004) between treatment groups and across sampling time points. In **A**, the letter represents results from a Tukey’s Honest Significant Difference test where groups that are connected by the sample letter are not statistically different. **D)** A dendrogram of the core microbiome based on sequence distances grouped taxa according to phylogenetic grouping. **E)** Bubble plots showing shifts in the relative abundance of the core members across treatment groups and sampling times points. Only a subset of the control samples collected at each sampling time point are shown under the ASW category. All treatment samples are displayed ordered by sampling time point and treatment conditions. The size of the bubble corresponds to relative abundance. If there is no point shown in the graph, the ASV was not detected in the sample. ASW = artificial seawater, FASW = filtered artificial seawater, ABS = Antibiotic solution. Results of statistical tests are reported in **Table S1** and the core community identified in **Table S3**.

We observed significant differences in alpha diversity during priming and recovery prior to and following antibiotic exposure (**Fig. 2A**). There was a significant interaction of treatment and sampling time (*p-value* < 0.0000367) on the overall number of ASVs per individual. The number of ASVs in the control samples did not significantly change over time (average nr. ASVs ± s.e. = 105.25 ± 0.6 to 75.8 ± 2.4). There was also no significant difference in number of ASVs in either of the ABS treatments during treatment (day 55, average nr. ASVs ± s.e. = 105 ± 6.1 to 86.6 ± 7.6). However, we did observe a significant decrease in the observed number of ASVs in anemones treated with ABS1 on day 61 and 76 (average nr. ASVs ± s.e. = 52 ± 13.9 to 51.8 ± 7.6) and anemones treated with ABS2 on day 76 (average nr. ASVs ± s.e. = 43.6 ±6.9). Indeed, the average number of ASVs per individual in anemones treated with ABS solutions was half that observed in the control samples (average nr. ASVs ± s.e. = 867.4±5.8) at the final sampling time point.

We performed a core microbiome analysis using anemones only exposed to ASW (control group). We chose to only include the control samples in our core analysis to determine how our experimental treatments impacted the composition of the core microbiome throughout the experiment. The core, as defined by taxa present in at least 90% of the control samples, consisted of 51 ASVs (**Fig. 2D**). The core ASVs included taxa belonging to the bacterial classes Actinomycetia (n=1), Bacteroidia (n=8), Polyangia (n=1), Phycisphaerae (n=1), Planctomycetes (n=3), Alphaproteobacteria (n=19), Gammaproteobacteria (n=15), Verrucomicrobia (n=1), and unclassified Proteobacteria (n=1; **Table S3**). The core community represents between 41 to 99% of each sample’s relative abundance (**Fig. 3**), indicating that we are capturing the majority of the community for most samples across treatment groups.

**Figure 3.**
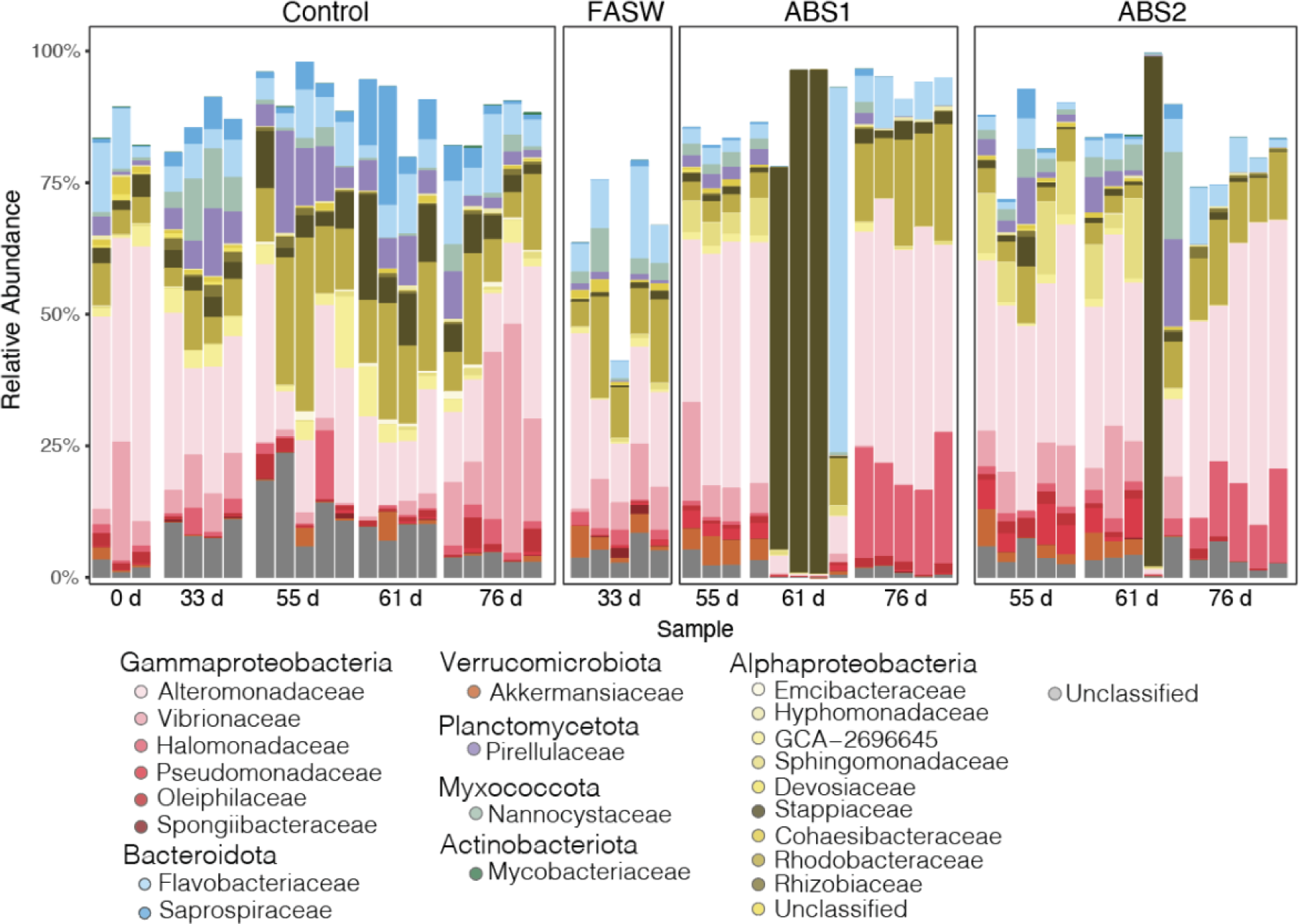
The Aiptasia core microbiome across treatment conditions. The bar plot revealed the relative abundance of each of the 51 ASVs that make up the core microbiome of the non-treated anemones. Members of the core microbiome were removed following exposure to FASW and both antibiotic solutions. At the final recovery time point (day 76), the microbiome became more consistent across individual anemones. ASVs are grouped and colored according to family level classification, each bar represents a single sample are grouped according to treatment and sampling time point prior to (0 d, 33 d) and directly following antibiotic exposure (55 d) and at both recovery time points (55 d, 76 d). ASW= Artificial Seawater, FASW = Filtered Artificial Seawater, ABS1=Antibiotic Solution 1, ABS2=Antibiotic Solution 2

Our results revealed that the core microbiome differed significantly following exposure to antibiotics. Using a PCoA analysis based on a Bray-Curtis distance matrix, we showed that the composition of the Aiptasia core shifted as a function of antibiotic exposure and sampling time point. A PERMANOVA test found a weak, but significant interaction of treatment and sampling time point on multivariate microbiome composition (adonis R^2^ = 0.14, p = 0.001; **Fig. 2C**). A subsequent beta-dispersion analysis showed that there were no significant differences between group dispersions (p-value = 0.271), or the distance to the group’s centroid. These results suggest that there is a strong separation in beta-diversity between treatments, with lowest beta-dispersion values for anemones treated with ABS1 solution sampled during the recovery time point (**Fig. 2B**).

Of the 51 core members of the Aiptasia microbiome, 31 were removed either directly after ABS exposure or were absent from the microbiome in treated anemones at either of the two recovery time points (**Fig. 2E**, **Table S3**). Of the seven taxa that were removed directly after ABS treatments, all ASVs, except for one species of Rhizobiales, remained knocked down during recovery. Knocked down taxa included the Actinomycetia species, *Dietzia psychralacliphila*, a species of Flavobacteriales*, Muricauda sp. 004804315,* the Gammaproteobacteria, *Spongiibacter tropicus*, and unknown species belonging to the Planctomycetes, Alphaproteobacteria and Gammaproteobacteria. Intriguingly, 23 of the 31 species that were ultimately removed from the Aiptasia core, were still present directly following ABS treatment. However, these taxa were lost or significantly reduced in the core community during recovery, which led to the ultimate reduction in bacterial diversity at the end of our experiment. Bacterial species that were removed included the Gammaproteobacteria *Oceanospirillum linum,* and the Alphaproteobacteria groups *Ruegeria sp.*, *Emicbacter sp.* and *Cohaesibacter sp*, and species within the Bacteroidia, Phycisphaerae, Planctomycetes, Alphaproteobacteria, Gammaproteobacteria and Verrucomicrobiales (**Fig. 2E, Table S3**).

### Differential patterns of microbial resistance, susceptibility, and selection by antibiotic exposure

Exposure to antibiotics differentially impacted the presence and absence of specific ASVs within the Aiptasia microbiome. We used the ALDEx2 package in R to first calculate the center log ratio to determine the within-sample, geometric mean of the read counts for each ASV (41, 42). We then used ALDEx2 to identify individual ASVs that were significantly different (Benjamini-Hochberg adjusted *p-value* < 0.05) between the ABS treatment and control group and had an ALDEx effect size greater than 2 or less than −2. For our analyses, we made two decisions to help guide the interpretation of our results. First, we chose to determine the differentially abundant (DA) ASVs for the entire dataset, rather than focus on the core community, which allowed us to identify all taxa that were impacted by the antibiotic exposure. We also chose to calculate the differential abundance of each ASV between the individual ABS treatment groups and the control at day 55, 61 and 76 to determine how each antibiotic treatment impacted the microbiome during treatment and recovery.

In total, we identified 37 unique ASVs that were differentially abundant across all comparisons (**Fig. 4**; **Fig. 5**). In our analyses, there were far more DA-ASVs identified during the final recovery time point (76 d) for both ABS1 (9 total; **Fig. 4C**) and ABS2 treatments (29 total; **Fig. 4C**). In contrast, we only observed four DA-ASVs in anemones collected directly after ABS1 exposure (**Fig. 4A**). We did not observe any DA-ASVs in anemones collected directly after ABS2 treatment (**Fig. 4D**). Of the differentially abundant ASVs, 65% were members of the Aiptasia core microbiome (**Table S4**).

**Figure 4.**
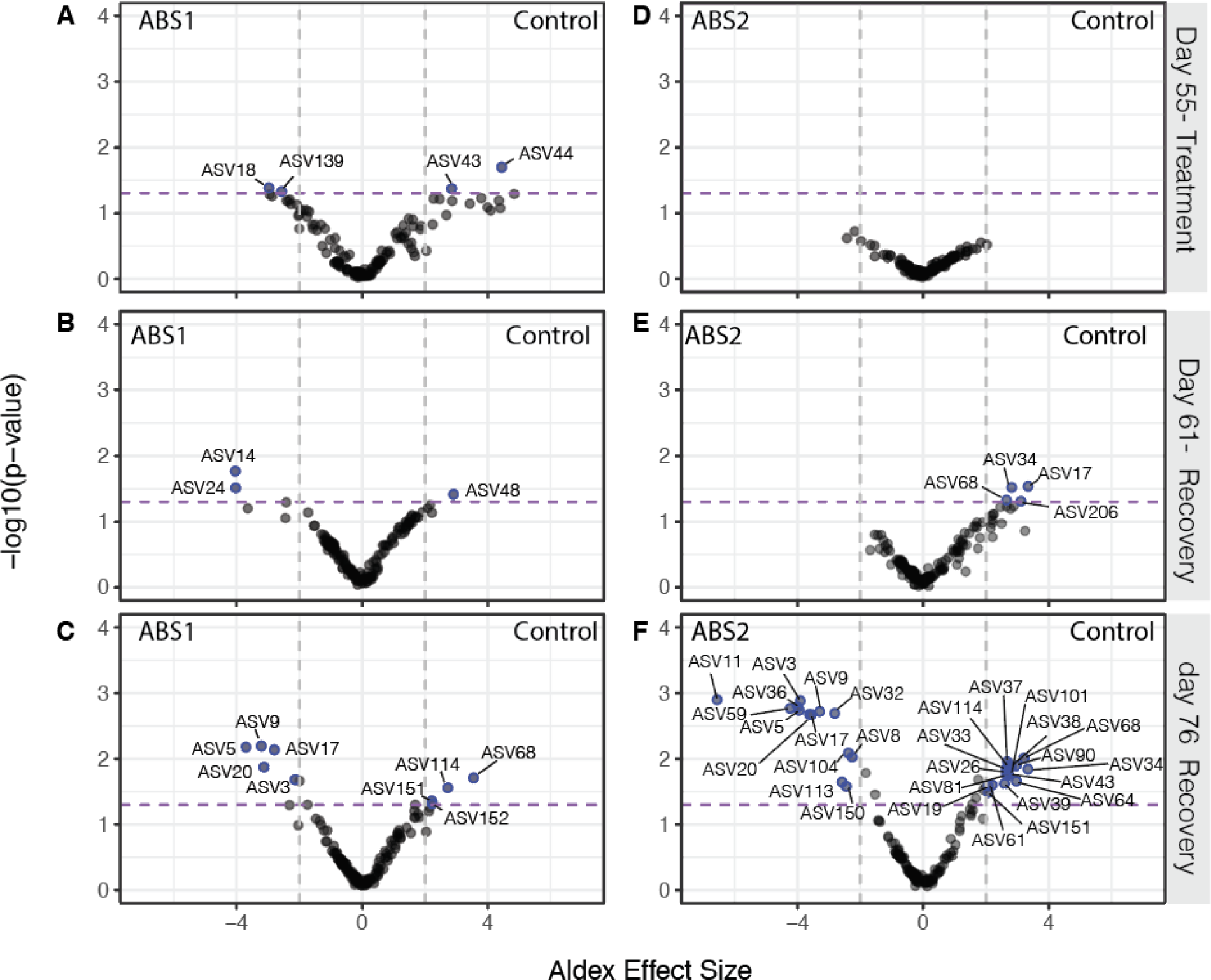
Exposure to antibiotics results in differential abundance of select ASVs 1 and 2 weeks after recovery. Volcano plot of all ASVs shows differentially abundant taxa (BH corrected T-Test p-value) between (**A-C**) ABS1 and control groups and (**D-F**) ABS2 and control groups (**A, F**) directly following ABS treatment (day 55), (**B,E**) one week post ABS treatment (day 61) and (**C,F**) three weeks (day 76) post ABS treatment. Taxa that are significantly different between treatment groups (Benjamini-Hochberg adjusted *p-values* < 0.05) and a calculated ALDEx Effect Size >2 or <-2 are represented by blue points and labeled with the ASV number. The purple dashed line represents a B-H adjusted, Welch’s t-test p-value < 0.05. The gray dotted lines are the threshold for the ALDEx Effect Size > 2 or < −2. ASVs that are enriched in the ABS treatments are plotted on the left side of the graph while ASVs enriched in the control are plotted on the right.

**Figure 5.**
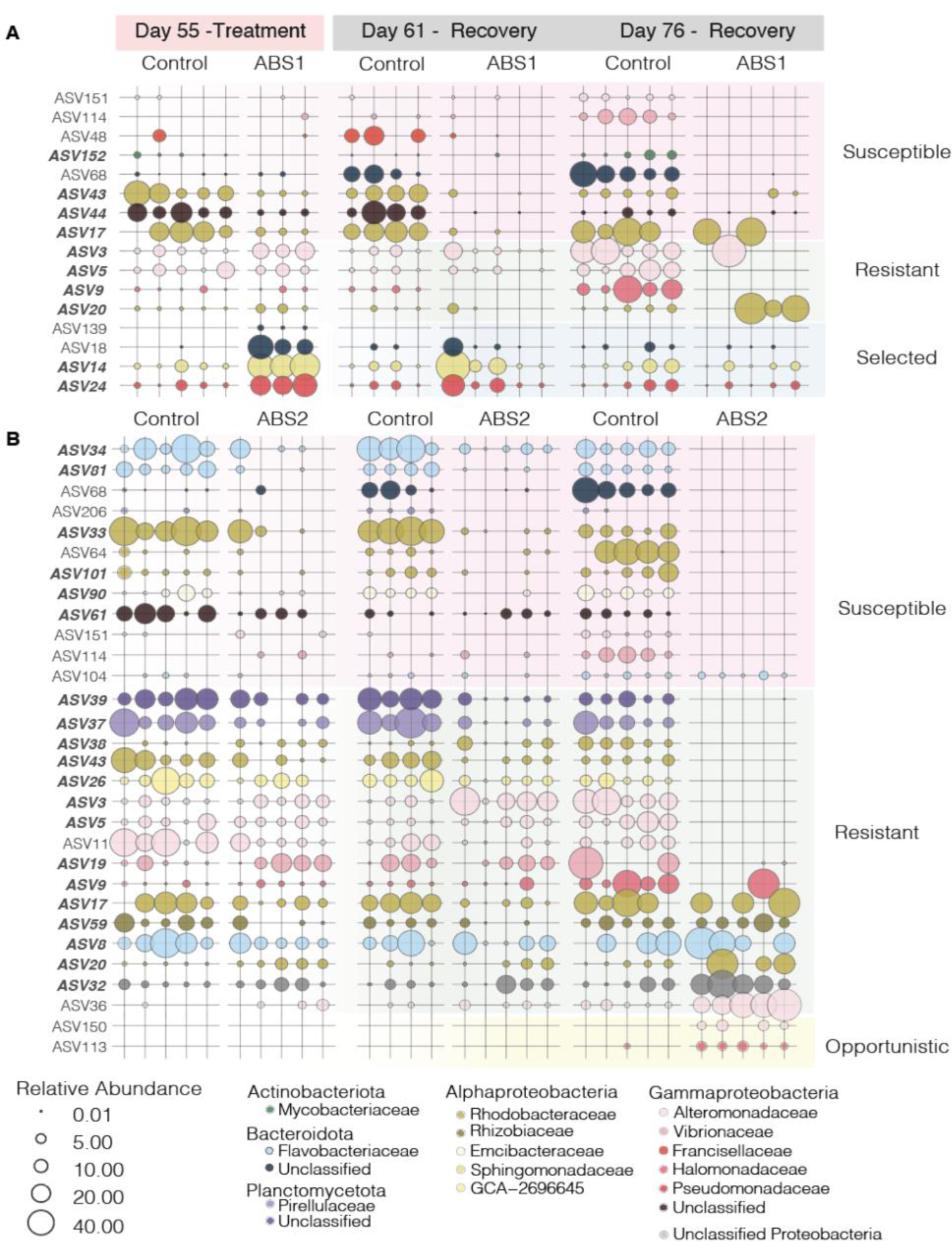
Relative abundance of differentially abundant taxa shows the ASVs were selectively knocked down during ABS exposure and remained depleted during recovery. The relative abundance of ASVs determined to be differentially abundant according to the ALDEx2 analysis (see Fig. S2) are plotted for (**A**) ABS1 and (**B**) ABS2. The size of each point represents the relative abundance of that taxa within the selected sample. Points are colored according to family level classification. ASVs lacking a point represent the absence of that ASV within the sample. Row shading is used to classify taxa based on their shifts in relative abundance throughout the experiment. Taxa were classified as susceptible or resistant to antibiotics, selected for by the antibiotics or were considered opportunistic taxa.

Next, we plotted the relative abundance of the 37 DA taxa across the three timepoints (day 55, 61, and 76) for both ABS treatments and control conditions (**Fig. 5**). To determine how each DA taxa is impacted by ABS exposure, we classified each ASV based on their shift in relative abundance directly after treatment and during recovery. ASVs were classified into one of four categories: (1) susceptible (2) resistant (3) selected and (4) opportunistic.

Susceptible taxa included ASVs that were either completely removed or significantly reduced in anemones treated with antibiotics directly following treatment (day 55). Susceptible taxa either remained reduced in relative abundance (e.g., ABS 44 in **Fig. 5A**), were eliminated during recovery (e.g., ASV 34 in **Fig. 5B**), or were removed during treatment but recovered to similar relative abundances as observed in the control group during recovery (e.g., ASV 104 in **Fig. 5B**). In total, we identified eight ASVs in ABS1 and 12 ASVs in ABS2 as susceptible to the antibiotics.

Taxa that were resistant to the antibiotics in the DA analysis included ASVs that had relative abundances at similar levels to the control group directly following antibiotic treatment, but either had significantly higher or lower relative abundances during recovery. For example, some taxa were not directly impacted by the ABS exposure during treatment, but during recovery they were removed from the microbiome (e.g., ASV5 in **Fig. 5A**). Conversely, other taxa that were not removed during exposure, ended up dominating the community after three-weeks of recovery (day 76) in FASW. Naturally, our DA analysis did not identify taxa that were resistant to ABS but did not change in relative abundance throughout the experiment (but see **Fig. 2**).

Finally, we also identified DA taxa that we considered to be selected for by the antibiotic exposure or were present in the microbiome as an opportunist. ASVs that had relative abundances that followed these patterns included taxa that were present in the microbiome of anemones that were treated with ABS solutions but were either lower in abundance or not present in the control samples. In total, we identified four taxa that were selected by the antibiotics in our ABS1 treatment. Selected taxa had higher relative abundances in individuals treated with ABS solutions in comparison to the control directly after treatment (55 d; **Fig 5A**), but the taxa were either removed (e.g., ASV139) or reduced to control levels during the recovery period (e.g., by 76 d). While there were no taxa that were selected by the ABS2 treatment, there were two taxa (ASV150 and ASV113) that we classified as opportunistic (**Fig. 5B**). These two taxa only appeared in anemones treated with ABS2 after three weeks of recovery in FASW.

## Discussion

### Antibiotic exposure reduced bacterial load up to one-week post treatment

In our study, we showed that antibiotic solutions decreased the microbial load and composition of the Aiptasia microbiome. Our primary finding is consistent with previous work in cnidarians which showed that antibiotics reduced the abundance of bacteria within the host microbiome following 5-7 days of exposure (31, 33, 43, 44). Surprisingly, our community composition analysis revealed that the microbiome only shifted in treated anemones during recovery. Indeed, by the final sampling time point (76 d), treated anemones were less diverse. We were also able to stably remove 29 of the 51 core members of the microbiome. Our results extend past studies seeking to manipulate and reduce the Aiptasia microbiome by demonstrating that it is indeed possible to use antibiotics to reduce the complexity of the community, maintain low microbial loads for up to one-week post treatment, and generate a new microbial stable state within the host.

In parallel to reducing the microbial load, we also observed that antibiotic exposure reduced host fitness, as measured by total biomass, *Symbiodinaceae* densities, and asexual reproduction rates. Because antibiotics are known to also impose metabolic consequences on eukaryotic cells (e.g., 42), it is challenging to definitively link reduction in biomass, asexual reproduction rate, and *Symbiodinaceae* densities to a reduction in microbial load. However, some interesting patterns emerged in our physiological data. Anemones treated with both ABS solutions overall had lower biomass throughout the experiment in comparison to the controls, except for anemones that had been treated with ABS2 where biomass seemed to recover to control levels at the final sampling time point. Apparent recovery of biomass could also be interpreted as a decrease in control anemone biomass due to reduced feeding at the end of the experiment. Another possibility is that anemones treated with both ABS solutions were unable to feed on sterile brine-shrimp throughout treatment leading to a reduction in overall biomass. This would match our observation that treated anemones did not expel visible food pellets following feeding. Intriguingly, Hydra are unable to feed when held in antibiotic solutions, however they do regain their feeding phenotype once transferred to fresh media (46). Our results suggest that antibiotic exposure in Aiptasia may have more serious phenotypic consequences then their freshwater relatives. Coupled to a reduction in biomass, anemones held in held in ABS solutions were not able to reproduce asexually during treatment. It is unlikely that the reduction in biomass alone led to a reduction in pedal laceration rates during antibiotic exposure as anemone growth and pedal laceration rates are not correlated (47–49). Furthermore, starved anemones tend to have increased asexual reproduction rates compared to fed individuals (47, 48). In the freshwater anemone, *Hydra*, antibiotic exposure also led to a reduction in anemone budding rates (50). Our result adds supports the hypothesis that the presence of bacterial partners plays a critical role in asexual reproduction of all anemones. Finally, our study revealed that antibiotic exposure also led to reduced *Symbiodiniaceae* densities in anemones treated with ABS1 solutions, but not with ABS2. Because we show differences in symbiont densities as a function of type of antibiotics indicates that either our ABS1 solution had a direct negative impact on the *Symbiodiniaceae* symbiont itself, or ABS1 differentially removes a key bacterial partner of the algae that is not targeted by ABS2 and resulted in the differential reduction of the algal population. Either way, our study extends past studies by quantifying the physiological impacts of antibiotic treatments on host fitness and suggests that bacterial load and composition may play an important role in the maintenance of Aiptasia fitness.

### Antibiotic exposure resulted in a less diverse, more consistent microbial community

Antibiotic exposure resulted in anemones with reduced complexity of the microbial community. Leveraging recommendations made by previous studies (31, 32), we successfully applied two different types of antibiotic solutions to individual anemones and observed a reduction in bacterial load and alpha diversity. While bacterial reduction only lasted one-week post ABS exposure, the composition was significantly shifted throughout recovery. We didn’t test the stability of the post-ABS community past three-weeks in FASW, but it is possible that the composition would have remained stable considering the anemones were held in sterile seawater and only fed antibiotic treated *Artemia* throughout the experiment. We suspected that we limited the introduction of new environmental bacteria to the host system, which enabled the long-term maintenance of the reduced microbial state.

Our work indicates that the recovery period following antibiotic exposure is critical for the establishment of a less complex microbiome model for Aiptasia. Recovery allows for the stabilization of the microbial communities and is likely impacted by the members of the microbiome that are differentially removed versus retained. Interestingly, not all cnidarians or marine invertebrates can maintain a low-diversity microbiome after antibiotic exposure. For example, the temperate coral *Astrangia poculata* recovered their microbial communities after only 2-weeks of recovery following exposure to antibiotics (33) and the Aiptasia strains, AIMS2, never showed a reduction in microbial diversity during or following antibiotic exposure (32). Similarly, antibiotic treatments, albeit much shorter than our three-week exposure, of the marine sponge, *Halichondria panciea*, resulted in a more diverse microbial community then control samples during recovery and ultimately lead to dysbiosis of the community (51). In *Acropora muricata*, bacterial communities failed to recover at least 4-days post antibiotic treatment (43).

Our work highlights the need for additional longitudinal studies showing the recovery dynamics of cnidarian microbiomes following antibiotic disturbance to better understand the role of individual taxa in microbiome recovery. While we were able to generate and maintain a lower diversity microbial community, we were not successful in completely removing the Aiptasia microbiome to generate axenic, or microbe-free, hosts. It is likely that our antibiotic treatments were not effective at reaching intracellular niches where microbial taxa might reside (e.g., those that live in symbiosis with *Symbiodiniaceae*) or that the Aiptasia host harbors antibiotic resistant bacteria that are challenging to remove with current drugs. One option, that until recently was not available as the life cycle of Aiptasia was not closed in the laboratory (52), is to apply antibiotics to newly settled Aiptasia larvae that lack their intracellular symbionts. Similar approaches that work to treat animals with antibiotics early in their life cycle have successfully resulted in axenic strains of *Drosophilia melanogaster*, *Caenorhabditis elegans*, and zebrafish (53–55), and could prove beneficial for generating germ-free cnidarians.

### Does Aiptasia microbiome have an alternative stable state following antibiotic exposure?

Our antibiotic treatments of Aiptasia resulted in an alternative stable state of the microbiome. The experimental approach transformed a flexible, dynamic microbial community into a microbiome that was narrow but more consistent across the population. For example, samples treated with antibiotics had a reduced population of ASVs belonging to Alteromonadaceae, Rhodobacteraceae, and Flavobacteriaceae families. These bacterial groups are commonly associated with the mucus microbiome of cnidarians, including a major component of the Aiptasia core microbiome (10, 56). Furthermore, species belonging to the Alteromonadaceae likely also play an important role in sulfur cycling by degrading DMSP (57). We also were able to remove members of the marine pathogen family, Vibrionaceae, suggesting that antibiotic treatments may help to restrict the growth of these potentially pathogenic microorganisms within the Aiptasia microbiome. The long-term removal of these bacteria species is promising for performing manipulative experiments to test the function of these species within the metaorganism. Conversely, some species of Halomonas, Pseudoalteromonas, Rhodobacteraceae, and Winogradskyella were either resistant to the antibiotics and were competitive dominates (e.g., ASV9, ASV17) or opportunistically colonized the host during recovery (e.g., ASV36, ASV113). The role and metabolisms of these ASVs within Aiptasia are currently unknown and remain targets for cultivation and cultivation-free analysis.

The establishment of alternate stable states during microbiome recovery from antibiotic treatments is well studied in vertebrate systems (58–60). Perhaps the best studied examples are from the human gut, where antibiotic treatments can lead to alternative stable states in microbiome populations, some of which are healthy and others diseased (61, 62). Furthermore, the recovery of the microbiome is often variable, incomplete (58–60), and can lead to the establishment of antibiotic resistant microbial species, which can make the host more susceptible to invasions by pathogens or subsequent exposure to stress (61, 62). In our study, we observed a consistent and stable recovery of the Aiptasia microbiome. The observed stability in microbiome recovery could be a result of the limited introduction of outside microorganisms by feeding the anemones with antibiotic-treated brine shrimp and holding them in sterilized artificial seawater, conditions that likely do not mimic recovery dynamics of similarly treated mammalian systems (e.g., 61, 62). Alternatively, the observed stability may indicate that the animal host has control over the proliferation, or lack thereof, of some microbial lineages after disturbance with the antibiotics. Further work should aim to test different genetic backgrounds of the animal host to explore if genotypic diversity may influence microbiome recovery trajectory in this emergent model system.

Determining factors that govern the susceptibility or resilience of cnidarians following microbiome manipulations is timely given the growing trend in studies looking to challenge, narrow or expand the associated microbial community (e.g., heat stress, antibiotic exposure, probiotics)(25, 31, 65). In our study, we showed that the Aiptasia system represents a powerful model for studying cnidarian resilience with shifting microbial diversity. Despite previous reports, it is possible to generate gnotobiotic individuals with a narrow and defined microbial community, without detrimental consequences to host health. However, it is unclear how the reduction of the microbiome will impact metaorganism response to subsequent disturbance events. Identifying the role of the complexity of the microbiome in cnidarian response to stress remains an essential priority for future studies.

## Materials and Methods

### Animal husbandry and experimental overview

We conducted a 76 day long experiment using *Exaiptasia diaphana* individuals, originally sourced from Hawaii (H2), to quantify the impacts of two different antibiotic solutions on the Aiptasia microbiome. The experiment was divided into three stages: priming (days 0-33), treatment (days 33 - 55) and recovery (days 55 - 76)(**Fig. 1A**). The priming stage enables the initial depletion of the Aiptasia microbiome (40), while exposure to each of the antibiotic solutions during the treatment phase helps to reduce the cnidarian microbial load (31, 32, 44, 50). Throughout the experiment, anemones were reared in 6-well plates at 25°C and 12 h light; 12 h dark cycle using 44 ± 15 µmol photon/m^2^/s light levels, in either artificial seawater (ASW; control group; Coral Pro Red Sea Salt Salinity 35 ppt) or 0.2 µm filtered artificial seawater (FASW; treatment groups). Anemones were either fed with freshly hatched non-treated (control group) or microbially depleted (treatment group) *A. nauplii* (31). To prevent the build-up of biofilms, water was exchanged from the welled plate after at least 4 hours after each feeding and anemones were transferred to fresh sterile plates once per week throughout the experiment.

Following the 33 d priming stage, anemones held in FASW were randomly divided into two groups and treated with either Antibiotic Solution 1 (ABS1; 50 µg / mL of rifampicin, chloramphenicol, nalidixic acid, and carbenicillin in FASW) or Antibiotic solution 2 (ABS2; 50 µg / mL of rifampicin, streptomycin, neomycin, and penicillin in FASW). We chose to use two different solutions in order to (1) determine if one solution is more effective than the other, (2) compare differences in microbial community composition as different types of antibiotics have varying mechanisms of action, and (3) because both solutions are known to deplete cnidarian microbiomes (31, 50). In preparation for treatment, mucus from individual anemones was removed via pipetting, after which the anemone was transferred to a sterile 6-well plate. Anemones were then treated with freshly made ABS1, ABS2 or ASW (control) solutions. During the 21 d treatment stage, treatment solutions were changed daily at the start of the dark cycle to avoid photodegradation of the rifampicin.

To determine how long treated anemones remained microbially depleted, we monitored the recovery of the Aiptasia microbiome for 21 days. Treated anemones were held in FASW. During the first two weeks of the recovery phase, the anemones were not fed to avoid introducing new microbial species to the previously depleted anemones. After which, they were fed microbially depleted *A. nauplii* twice per week.

### Anemone sampling

To quantify the impact of ABS treatment on the anemone microbiome, we sampled replicate anemones on day 0 (baseline; n=3), 33 (priming; n=5), 55 (treatment; n=5), 61 and 76 (recovery, n=5). To avoid sampling the microbiome associated with the ingestion and digestion of the *A. nauplii,* all anemones were sampled at least four days following the previous feeding day (Fig 1A). Prior to sampling, anemones treated with ABS solutions were transferred to FASW for 24 hours. Individuals were then homogenized in 250 µl of FASW using a pestle motor. An additional 300 µl of FASW was added to each sample to bring the total volume of anemone homogenate to 550 µL. The resulting homogenate was mixed and separated into individual aliquots in preparation for quantifying bacterial load (50 µL), determining algal symbiont densities and protein content (180 µl with 20 µL of 0.1% Sodium Dodecyl Sulfate [SDS]), extracting DNA (250 µL). The homogenate sample prepared for DNA extraction was transferred to a bead beating tube (1:2 mixture lysing matrix B:D MP Biomedicals) containing 500 µl DNA/RNA shield (Zymo Research) and subsequently cells were lysed using a Fastprep 24-5G (MP-Biomedicals; 2 cycles of 8m/s for 60 s with a 5 min pause between cycles). All aliquots of the original homogenate, except that used to determine microbial load, were held at −80°C until further analysis. Finally, at each timepoint, we also sampled 1 L of FASW by collecting filtrate on a 0.22 µm filter cartridge (Millipore-Sigma Cat # SVGP01050) to quantify the composition of the microbial communities within the FASW. After filtration, the cartridge ends were sealed and stored at −80°C until DNA extraction.

### Bacterial Load

We quantified the bacterial load from individual anemones using a colony forming unit (CFU) assay. Briefly, 50 µL of the anemone’s homogenate was serially diluted 10-, 100-, 1000- and 10000 fold and 50 µL of each dilution was plated on Marine Agar plates (Difco Marine Broth 2216 with agar). Plates were incubated at 28℃ for at least 24 hours and up to 2 days, after which colonies were counted. We compared CFU counts between treatment groups at each sampling time point independently using a Kruskall-Wallis nonparametric test to determine the effect of treatment (ABS1, ABS2, ASW) on the number of CFUs / mL.

### Microbiome Data Collection

In preparation for DNA extraction, the lysed anemone homogenate was defrosted on ice. Filter cartridges were defrosted on ice and DNA/RNA shield was added directly to the filter column at the start of defrosting, then drained into the bead beating tube. Filters were aseptically removed from the column and sliced into strips before being placed in the bead beating tube and homogenized as above. Zymobiomics DNA miniprep kit (Zymo Research) was used to extract DNA from all samples following the manufacturer’s protocol with the following modification: DNA was eluted from the binding column in two separate, 1 min incubations using 50 µL of nuclease free water to result in final volume of 100 µL. Resulting nucleic acid concentrations were measured using the Qubit dsDNA (double-stranded DNA) broad range assay kit (Invitrogen) on a Qubit 4 Fluorometer (Invitrogen).

To generate the sequencing library targeting the V4 region of 16S rRNA gene, DNA extracts were amplified in duplicate reactions using the barcoded primer set 515FY, 806RB (66, 67). Briefly, PCR reactions were composed of 2 - 4 µl of DNA template, 1.25 µl of each primer, 12.5 µl of Q5 High Fidelity 2X Master Mix (NEB), and 7 - 9 µl of nuclease free water for a final volume of 25 µL. The amount of DNA template added to each reaction was dependent on the amount of template needed to achieve amplification. PCR reactions were carried out using a BioRad C1000 thermal cycler (BioRad Laboratories) with the following gradient: an initial denaturation step at 98℃ for two minutes followed by 30-38 cycles of denaturation at 98℃ for 20 sec, 55℃ for 15 sec, 72℃ for 3 minutes, and a final extension of 72°C for five minutes. All reactions were held at 4°C. Duplicate PCR products that successfully amplified were pooled. 24 µL of the pooled reactions were visualized on a 1 % agarose/TAE gel. Using a Quick-Load Purple 100bp DNA Ladder (NEB) standard, the band corresponding to the expected size of the V4 region of the 16S rRNA gene was excised from the gel and extracted using the Monarch DNA Gel Extraction Kit (NEB). Purified PCR products were pooled to equal molar ratio (2 ng/ µL) and submitted to University of California, Davis’ Genome Center for sequencing using 250-bp paired-end MiSeq (Illumina) (9).

### Microbiome Analysis

Demultiplexed fastq files were processed in R Studio(v. 1.1.463) using the DADA2 pipeline (v.1.24.0)(66) and filtered using the FilterandTrim function using default parameters with the following exceptions: TruncLen = c(150,150), maxEE=c(2,2), and multithread=True. Sequences were subsequently denoised, merged and chimeras were removed, bringing the total number of sequences in the data set from 10,796,554 to 9,768,428. Samples with less than 20,000 reads or greater than 250,000 reads were removed from the dataset. 3012 ASVs were identified in the dataset and taxonomy was assigned using the Genome Taxonomy Database R.202 (69). Assigned Taxa, ASVs, and sample data were merged into a single phyloseq object for subsequent analysis (70). After which, ASVs with taxonomic strings matching chloroplasts, mitochondria, and eukaryotic identifications were removed from the dataset resulting in 2800 ASVs. A multiple sequence alignment was run using the AlignSeq function from the DECIPHER package (71), and a neighbor joining tree was created and fitted for maximum likelihood using a GTR model with the phangorn package (72). The resulting tree was added into the phyloseq object. In order to remove contaminating sequences from our phyloseq object, we used the decontam package to help identify ASVs that are not part of the Aiptasia microbiome using two steps (73). First, using the isContaminant function, we identified 28 ASVs as kit contaminants by applying a prevalence threshold of 0.1 to the extraction blanks. Additionally, we removed 2280 ASVs with only one occurrence in the dataset. Because we also wanted to remove ASVs that were primarily within the filtered ASW, we then performed a second decontamination step that leveraged the seawater blank samples to filter out an additional 80 ASVs based on the same prevalence criteria. After removing the contaminating sequences, we removed all extraction and seawater blanks from the dataset which resulted in a phyloseq object of 82 samples composed of 412 ASVs. The full phyloseq object was used for microbial diversity analysis described below.

First, to determine the impact of antibiotics on bacterial alpha diversity we used phyloseq to calculate the observed number of ASVs per sample using the entire 412 ASV data matrix. We then calculated a two-way ANOVA to quantify shifts in the total number of ASVs across sampling timepoints and treatment groups. Differences in treatment levels were determined using a Tukeys HSD test.

Next, we chose to quantify shifts in sample beta-diversity of the Aiptasia core microbiome. To calculate the core, we first filtered the phyloseq object to only include samples held in ASW (control) and ASVs with non-zero counts. We then determined the core microbiome using the *core* function within the microbiome package (v1.22.0)(74) by retaining ASVs based on prevalence (i.e., ASVs present >90% of control samples). After normalizing ASV counts to relative abundance within the original phyloseq object, we filtered the ASVs to only include the core community. The resulting phyloseq object was subsequently used to calculate a Bray-Curtis distance matrix that was used to inform a Principal Coordinate Analysis (PCoA). Differences between sample groups were tested using a permutation multivariate ANOVA (PERMANOVA) using the *adonis2* function within vegan (v.2.6-4)(75). We also calculated and compared the distance to the group centroid using the *betadisper* function in order to determine within group variance of the microbiome.

Finally, to determine which taxa are differentially enriched directly after treatment (55 d) and during recovery (61 d and 76 d), we used the Aldex2 package (v.1.28.1)(41) to compare variation in the abundance of ASVs between the control and each treatment group at each sampling time point. We chose to use the entire datasets for the differential abundance analysis to ensure we were not missing key taxa that were not part of the core that either significantly increased or decreased in abundance during the experiment. Taxa were determined to be significant if they had an effect size > 2 and a *p-value* < 0.05 for both aldex.ttest and aldex.glm at any of the three timepoints for either the control vs. ABS1 and control vs. ABS2 comparisons. After determining which taxa were differentially abundant, we plotted relative abundances across treatment groups and time points (55 d, 61 d and 76 d) and classified individual taxa into groups based on their shift in relative abundance directly after treatment and during recovery.

### Aiptasia physiology

To quantify the impacts of the antibiotics on host metaorganism physiology, we monitored shifts in total protein, algal densities, and asexual reproduction rates throughout the experiment. To assess shifts in biomass, anemone homogenate was defrosted and 20 µl was used in a Pierce BCA protein assay to quantify total protein in triplicate according to manufacturer instructions (Thermo Scientific). To quantify algal cell density, we defrosted 180 µL of the anemone homogenate reserved for algal cell counts and diluted the homogenate 1:20 in FASW containing 0.01% SDS. Animal tissue cells in the diluted surrey were sheared with a 25G needle. Algal cell counts were obtained from 100 µL of each sample using flow cytometry (BioRad ZE5 Cell analyzer) using the following parameters: 405 nm laser forward scatter, 448 nm laser side scatter, and Chlorophyll fluorescence with a 448 nm laser excitation and detection with a 692/80 bandpass filter. The plate was agitated every 5 samples and ran at a flow rate of 1.5µL/sec and resulting cell counts were normalized to total protein content (76). To quantify the rate of asexual reproduction, we held a separate set of replicate (n=6) anemones in six well plates and treated them as described above for the experimental anemones. Throughout the experiment, we counted the number of produced pedal lacerates on a weekly basis. Counted pedal lacerates were removed from each well following counts. All metrics were compared across treatment groups and sampling times points after being checked for normality using a Shapiro-Wilk test. Non-normal datasets (CFU counts and pedal lacerates) used a Kruskal-Wallis test with Dunn’s post-hoc comparison, while normally distributed parameters (protein content and log-adjusted symbiont density) used an ANOVA with Tukey’s Honest Significant Difference using the stats package in R.

## Supplemental Tables

**Table S1.** Model statistics for treatment comparisons across physiological and microbiome variables.

**Table S2.** Means and standard error for physiological variables across experimental treatments.

**Table S3.** Members of the Aiptasia core microbiome.

**Table S4.** Differentially abundant taxa and patterns of removal.

## Data Availability

Raw sequences are available at the National Center for Biotechnology Information’s Sequence Read Archive under PRJNA1048100. Physiological data presented in this manuscript can be found at https://doi.org/10.5281/zenodo.10257452. Code is available at https://github.com/smacvittie/aiptasia_abs_recovery.

## Acknowledgements

We would like to thank the UC Merced Stem Cell Instrumentation Foundry and Dr. David Gravano for assistance in generating flow cytometry data. Research was supported in part by the DoD Research and Education Program for HBCU/MSI Instrumentation Grant W911NF1910529. The sequencing was carried out at the UC Davis Genome Center DNA Technologies and Expression Analysis Core, supported by NIH Shared Instrumentation Grant 1S10OD010786-01. The authors thank Dr. Anya Brown for fruitful discussions related to the manuscript. Funding from a National Science Foundation Grant to E.M.S. (NSF BRC-Bio #2217769) provided financial support for this work.

